# A global P-Process map to avoid P-value abuse

**DOI:** 10.1101/2025.04.06.646254

**Authors:** Jing Xu, Siyu Wei, Junxian Tao, Chen Sun, Haiyan Chen, Lian Duan, Zhenwei Shang, Wenhua Lyu, Hongchao Lyu, Mingming Zhang, Yongshuai Jiang

**Author notes:** Correspondence*: Yongshuai Jiang, College of Bioinformatics Science and Technology, Harbin Medical University, 194 Xuefu Road, Nangang District, Harbin, Heilongjiang Province, China. These authors contributed equally to this work (Jing Xu, Siyu Wei, Junxian Tao, Chen Sun).

## Abstract

P-abuse is serious in difference identification analysis of data. How to avoid P-abuse is a huge challenge. Here, we evolve P-value (a single value) to P-Process (a global landscape of P-values under different sample size) to help researchers correctly recognize and use p-value. We observed -ln(P-Process) after rotation has very similar morphology with Wiener Process (or Brownian motion). Based on this property, for any sample size, we estimated the 95% fluctuation range of P-value and further estimated how many samples (*N*_*95*_(*α*)) could make sure 95% of P-values less than the given significant level *α*. Tests proved that the estimation obtains a good performance with only a small number of samples in each group. For broader accessibility, a free web-service, P-Process Map, is available online to show the whole landscape of P-Process. At the end of the article, we explained 10 of the most typical P-abuse problems which can be easily voided by using P-Process. This “rethinking” of P-value from a higher position, which is profoundly different from the way we have seen this for the past century, would yield a new era - 2P era in which hypothesis testing is strongly required to evolve from P-value to P-Process.

## 1. INTRODUCTION

Fishing for difference between two groups plays increasingly important roles in various fields. Still hypothesis test with its arbitrary P-values is the main approach for difference identification(Goodman, 1993). However, P-value, the “gold standard” of statistical validity, is not so reliable as many scientists assume. There are serious problems about the use of P-value(Fidler, Burgman, et al., 2006, Hoekstra, Finch, et al., 2006, Lena and Hipp, 2014, Schatz, Jay, et al., 2005). An analysis of 406 articles published in the Archives of Clinical Neuropsychology found that 67.6% of them were revealed inappropriate use of P-value(Schatz, Jay, et al., 2005).

The most common problem is claiming there is “no difference” between two groups just because the P-value is larger than the threshold (usually be 0.05) specified in advance(Goodman, 2008, Goodman, 1989, Halsey, Curran-Everett, et al., 2015). Surveys of 791 articles have found nearly half of them mistakenly use P-value like this(Amrhein, Greenland, et al., 2019), which has greatly contributed to the publication bias. There are many other P-value misconceptions(Goodman, 2008) such as talking about the P-value without considering sample size, concluding that two studies conflict because different P-values are obtained(Camerer, Dreber, et al., 2016, Camerer, Dreber, et al., 2018), making statistically significant P-value represent clinically important effects, granting the null hypothesis a chance of being true by the P-value, etc.

These widespread abuses of the P-value have largely contributed to increasing unrepeatable scientific results. It turned out that these problems were not only caused by data or analysis but largely by the surprising slippery nature of P-values(Nosek, Spies, et al., 2012, Nuzzo, 2014) and people’s limited cognition about P-values. It is high time to go beyond the shibboleth and think about P-value under a higher level.

## 2. P-value Not Significant Enough Declares No Difference? Absolutely Not!

P-value is widely used for difference analysis, but there is a little-known fact among non-statisticians that P-value gradually decreases with increasing sample size majority of the time(Albers, 2019, Demidenko, 2016, Visscher, Wray, et al., 2017). So it would be unwise to draw firm conclusions whether there is difference between two groups from the single P-value of only one hypothesis test(Goodman, 2008). If your P-value is not significant enough, do not give up, it doesn’t necessarily mean no difference but may remind insufficient evidence for difference (see SOM Section 2). Our results also confirmed that.

We have explored two continuous normally distributed populations, A and B, with different mean values (0 and 1) and the same variance (3). Considering the balance of sample sizes in the two groups, we drew *n* samples from A and B separately, generating two datasets S1 and S2. By exploring *n* from 2 up to 250, results showed that P-values of t test became as small as desired by increasing the sample size, and the differences observed were clearer (Fig.1 A, B). An example of differentiating gender in the real world illustrated this phenomenon well. By analogy with sample size as resolution, the higher the resolution, the more the information stored in the image and the clearer the painting. Likewise, the larger the sample size, the more significant the differences (Fig.1 C).

**Fig. 1.**
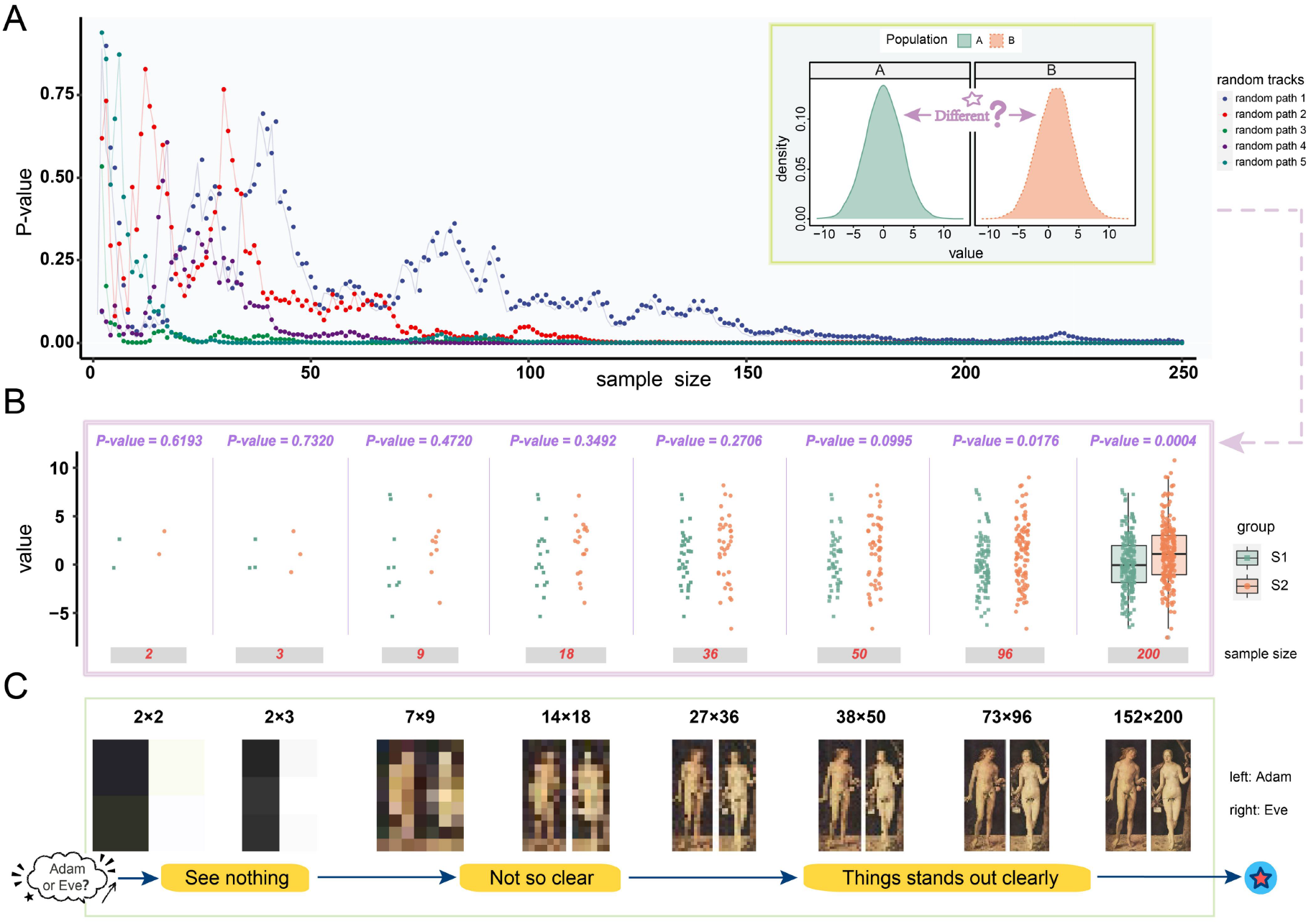
P-values decrease with sample size increases. **(A)** Scatter plot for P-values of t test carried on the two normally distributed population A and B, with different mean values (0 and 1) and the same variance (3). The sample size was explored from 2 to 250. The process was randomly repeated for five times. **(B)** Example from (A), random path 2. The datasets selected from A and B were drawn with dots of different shapes and colors. On the bottom are sample sizes, on the top are corresponding P-values. **(C)** “Adam and Eve” (World famous oil painting by Albrecht Durer) seen at different resolutions.

The similar feature was discovered in other hypothesis test methods (see SOM Section 3). Here we chose t test as an example for demonstrating during the subsequent sections.

## 3. P-Process

Here, we demonstrate that P-value of hypothesis test for difference analysis is not a single constant value, but in fact a quasi-stochastic process (Fig.2 A), referred as P-Process and defined as {*P*(*n*), *n* ∈(*0*, ∞)}. P-Process shows a global landscape of P-values under any sample size. The fact behind P-Process is that the uncertainty about difference gradually decreases (information obtained increases) with sample size growing and the difference gradually emerges and becomes clear. This well reflects people’s recognition process of difference, so that P-Process can be also described as Difference Recognition Process. Getting the description of two attributes for P-Process (geometrical mean and 95% fluctuation range), would be a great help for further/deeper understanding and better use of P-value.

**Fig. 2.**
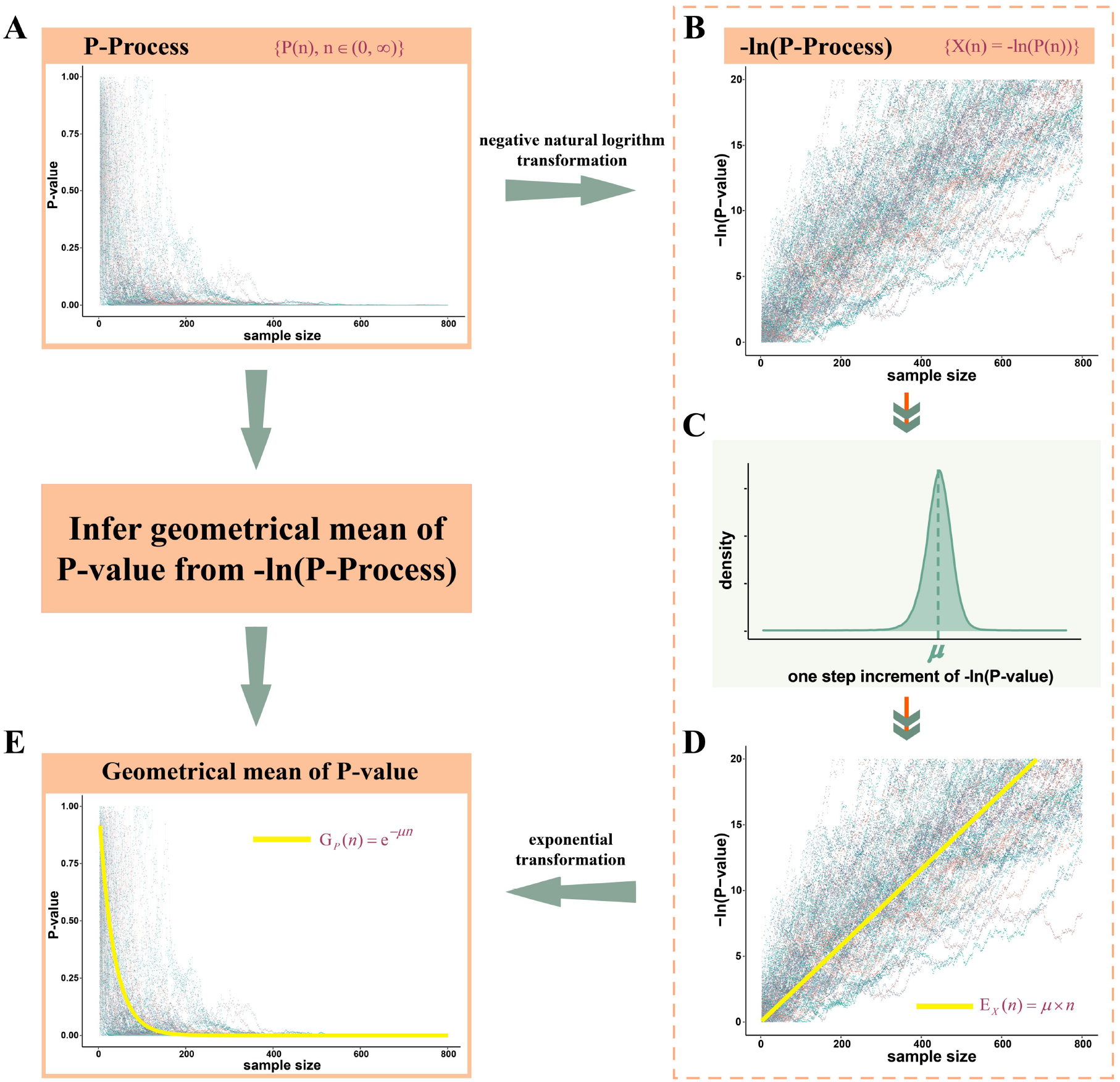
Flowchart for inferring geometrical mean P-value from -ln(P-Process). 100 paths of **(A)** P-Process and **(B)** -ln(P-Process) for two binomial datasets *N*(0, 3) and *N*(1, 3). **(C)** Distribution for one step increment of -ln(P-value). **(D)** Mean of -ln(P-value) under any sample size *n*: *EX*(*n*). **(E)** Geometrical mean of P-value under any sample size *n*: *GP*(*n*). The yellow line represents the geometrical mean of P-Process.

### 3.1 Infer geometrical mean of P-Process

Negative natural logarithm of P-Process, {*X*(*n*) = − ln(*P*(*n*)), *n* ∈(*0*, ∞)}, is also a stochastic process, referred as -ln(P-Process) (Fig.2 B). We observed that there exists a good linear correlation between -ln(P-Process) and sample size, stemming from the increasing amount of information we obtained. We firstly calculated the average increment of -ln(P-value) for each sample increase, referred as *μ* (Fig.2 C). With *μ* for the average growth trend of -ln(P-Process), then for any sample size, the mean value for it was estimated with a simple linear function *E*_*x*_(*n*) = *μ* × *n* (Fig.2 D, SOM Section 4.3). This enables further estimation for geometrical mean of P-values under any sample size, which can be calculated as *G*_*p*_(*n*) = *e*^−*μn*^ (Fig.2 E, SOM Section 4.4).

### 3.2 Infer 95% fluctuation range of P-Process

Obviously, P-value under certain sample size fluctuates at different paths because the samples were randomly selected (Fig.3 A). And having only geometrical mean of P-values is often not enough to draw definitive conclusions about the difference, further information of fluctuation range is needed. We observed that the scale normalized -ln(P-Process), called *X* ^*unif*^ (*n*), after rotating 45 degrees clockwise, has similar morphology with Wiener Process {*W*(*t*), *t* ∈(*0*, ∞)}. The increment of *W*(*t*) is in normal distribution *N*(*0*, (*k*σ)^*2*^), where σ is the standard variation of one step increment for *X* ^*unif*^ (*n*). (Fig.3 B, SOM Section 4.5.1).

**Fig. 3.**
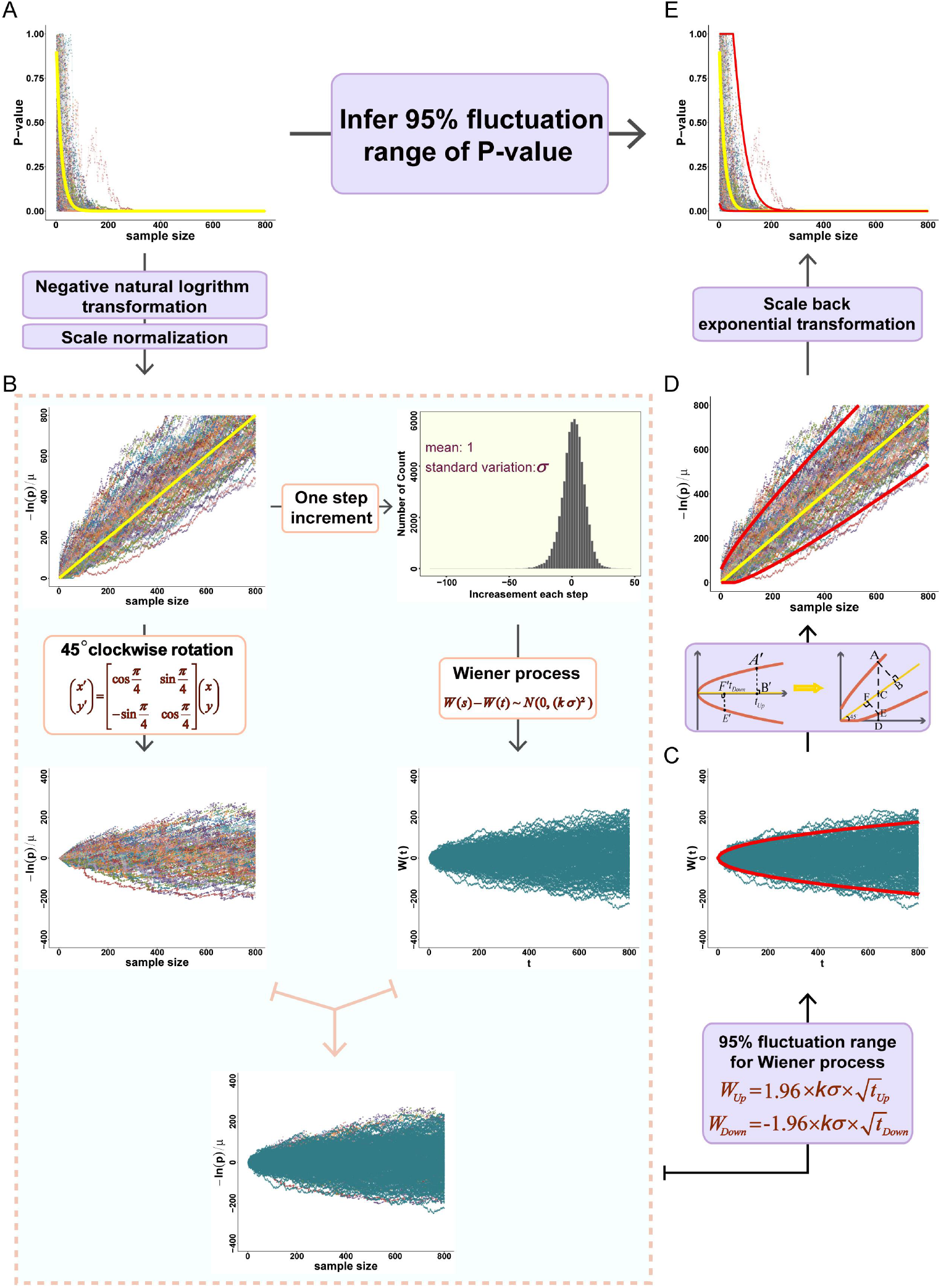
Flowchart for inferring 95% fluctuation range of P-value. **(A)** 100 paths of P-Process for two binomial datasets *N*(0, 3) and *N*(1, 3). **(B)** the 100 paths of *X*^*unif*^(*n*) after 45 degrees clockwise rotation significantly overlaps with the image of Wiener Process *W*(*t*). The 95% fluctuation range of **(C)** *W*(*t*), **(D)** *X*^*unif*^(*n*) and **(E)** P-Process. The red lines represent the fluctuation range.

We can easily calculate the 95% fluctuation range of *W*(*t*) on the basis of its variation formula (Fig.3 C):

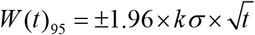

Here, *W*(*t*)_*95*_ represents 95% fluctuation range of *W*(*t*) and *t* represents the passing time. Each passing time *t* in *W*(*t*) potentially corresponds to a sample size *n* in *P*(*n*). Based on morphological similarity, the 95% range of fluctuation for *W*(*t*) can be used in place of that for *X* ^*unif*^ (*n*) (Fig.3 D, SOM Section 4.5.2):

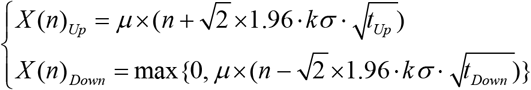

And then the 95% fluctuation range of P-Process can be estimated as (Fig.3 E):

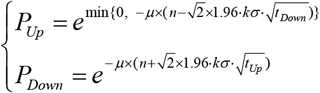

Here, *P*_*Up*_ and *P*_*Down*_ are respectively the upper and lower limits of fluctuation for P-value under the given sample size *n*, *t*_*Up*_ and *t*_*Down*_ are respectively the corresponding passing time along the mean direction in *W*(*t*) (SOM Section 4.5.2).

## 4. P-Process Map

To build a global view of P-value for better understanding of difference, we constructed a P-Process Map based on the properties of P-Process. P-Process Map shows (i) the paths of P-Process, (ii) the geometrical mean P-value under any sample size, (iii) the 95% fluctuation range of P-values under any sample size, (iv) the trend that P-value decreases and converges as sample size increasing to an infinitely large quantity (i.e., the trend that difference becomes visible) (Fig.4).

**Fig. 4.**
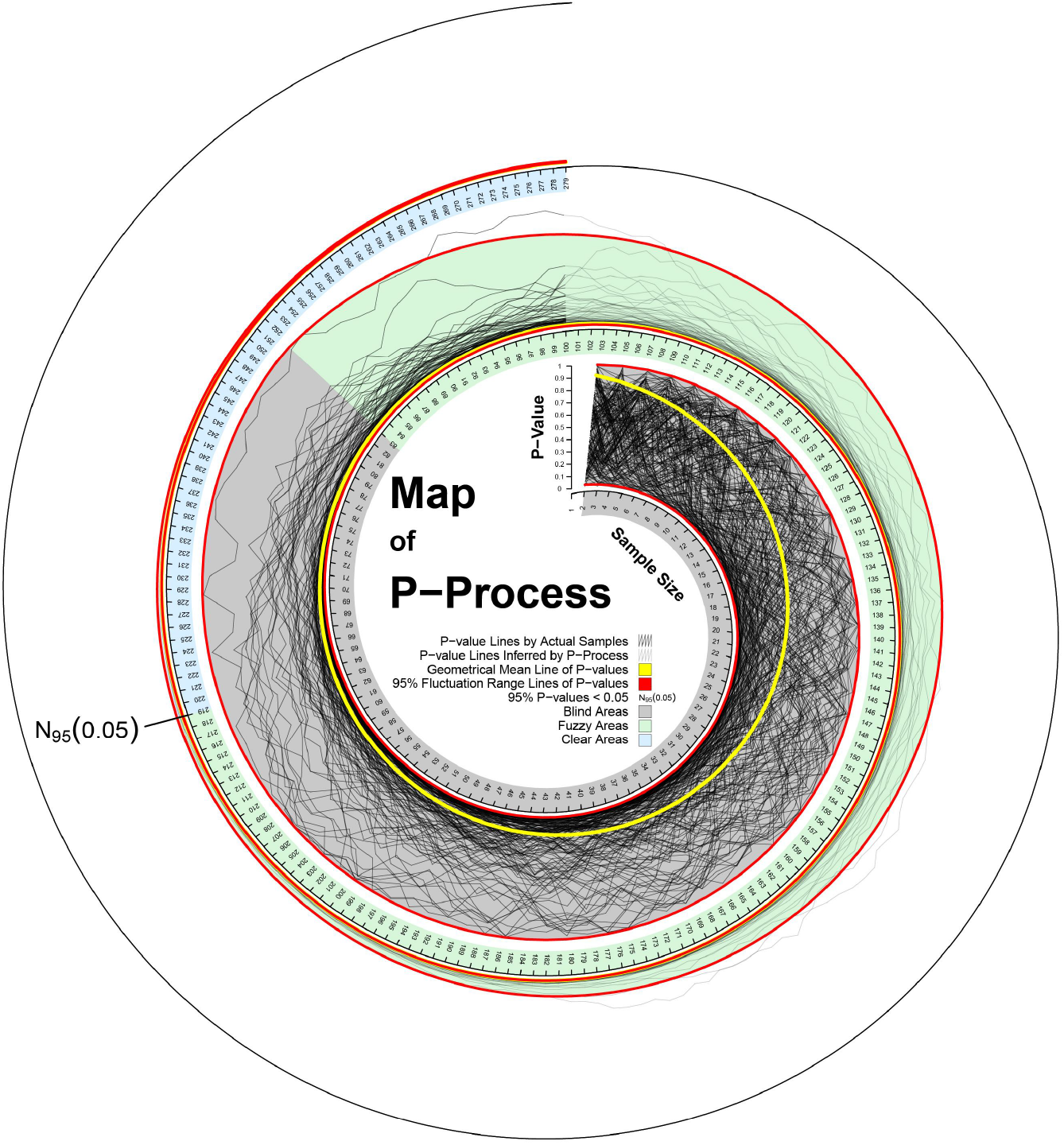
P-Process Map. The x-axis is the sample size in each group and the y-axis is the P-value. The dark black lines are those connected to the P-values calculated from the real data, the light black lines are those connected to the P-values inferred by the P-Process. The yellow line represents the geometrical mean, the red lines and the area enclosed by them represent the fluctuation range. The background color in gray is the blind area, green is the fuzzy area and blue is the clear area.

### 4.1 Three phases in P-Process Map

For better understanding and more intuitive description, P-Process Map is divided into three phases according to the upper 95% fluctuation range and a specific threshold *α* (such as 0.05): (Phase I) blind area, the upper bound of 95% fluctuation range stabilizes at 1 in this area, that is the uncertainty of difference won’t be eliminated at this stage even with sample size increasing; (Phase II) fuzzy area, the upper bound of 95% fluctuation range is between *α* and 1, which shows that the uncertainty of difference is constantly removed as sample size grows; (Phase III) clear area, the upper bound of 95% fluctuation range are all less than *α*, where difference stands out clearly even no sample added (Fig.4).

### 4.2 *N* _*95*_(*α*)

From P-Process Map, the minimal sample size required to make P-Process reach “clear area” can be defined as *N*_*95*_(*α*). *N*_*95*_(*α*) marks a turning point when difference can be clearly seen among 95% cases in difference recognition process.

Intuitively, various P-Process Maps are available for different difference sizes. A “Large” difference just goes through a short blind area and fuzzy area (Fig.5 A), that is only a small number of samples (*N*_*95*_(*α*) is small) are required to see this difference clearly. Conversely, it takes a long long period of blind area and fuzzy area before a “Small” difference coming out clearly (reach clear area) (Fig.5 B-E). Briefly, the larger the difference, the less the samples needed for recognizing the difference, the smaller the *N*_*95*_(*α*). We strongly recommend that researches provide *N*_*95*_(*α*) in difference analysis for convenient comparison of results in different situations.

**Fig. 5.**
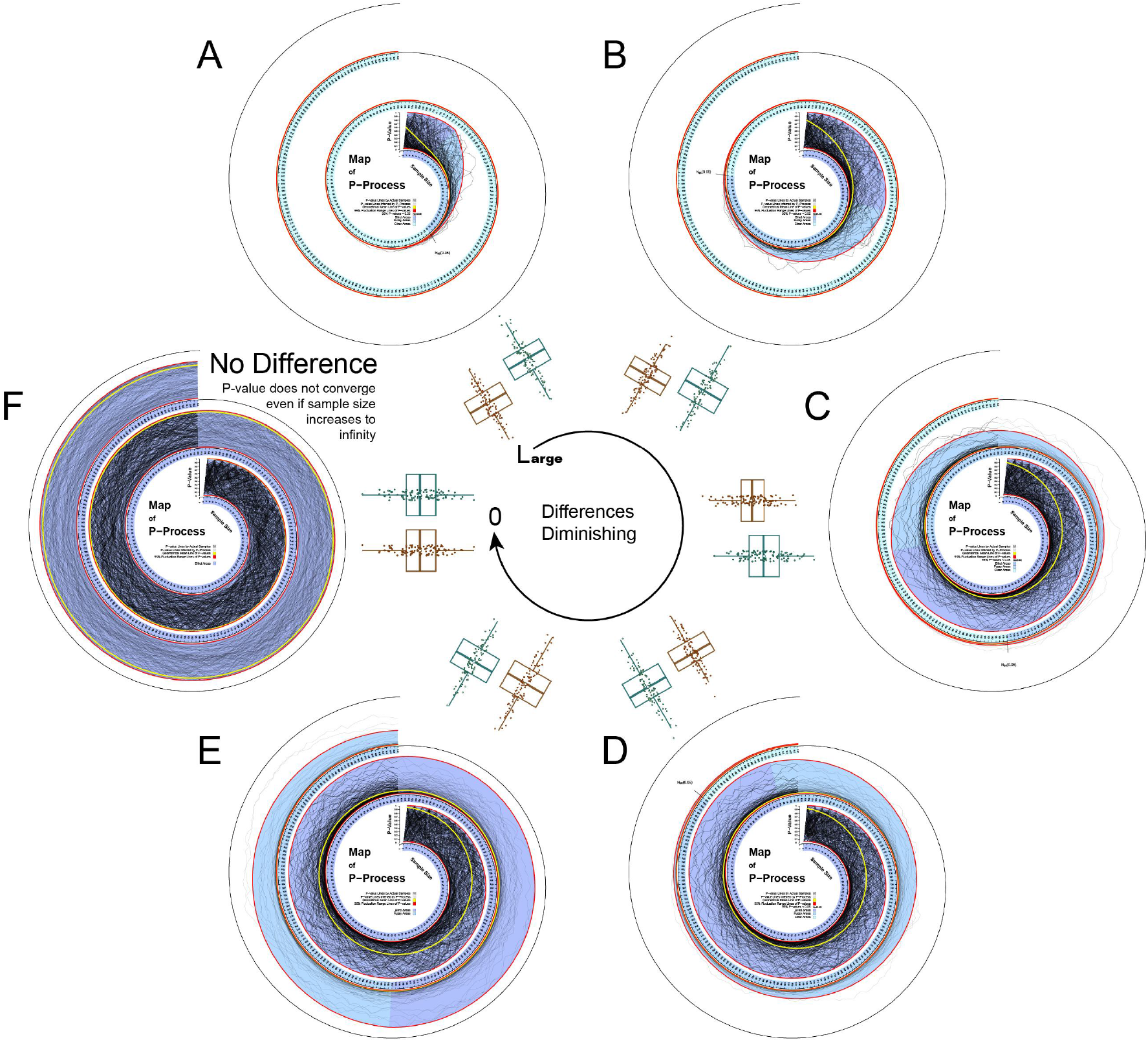
P-Process Map of difference with different sizes. P-Process of two normal populations with the same variation which is 3 and different mean values: **(A)** mean difference is 3, **(B)** mean difference is 2, **(C)** mean difference is 1.5, **(D)** mean difference is 1, **(E)** mean difference is 0.8, **(F)** mean difference is 0.

### 4.3 Note

As seen in the P-Process Map, for the most situations in the real world, all unbiased sample sets (X1 and X2) from two different populations (S1 and S2, E(X1) = E(S1), E(X2) = E(S2)) will eventually obtain a P-value small enough upon ever-growing sample size. The larger the difference between S1 and S2, the faster the P-value decreases and converges (Fig.5 A-E).

However, there is a circumstance that P-value will experience a particularly long period of blind area and never get to the clear area even if the sample size increases to infinity (Fig.5 F), which is what people usually consider as “no difference”. This is an extreme case of difference and is referred to as null hypothesis (H0: there is no difference between two groups) in hypothesis test of difference analysis, i.e., “the two datasets” studied were from the same population, (E(S1) = E(S2)).

## 5. Parameter stability

We tested the reliability of P-Process based on its two key parameters *μ* and σ, which are respectively for estimating geometrical mean and for inferring 95% fluctuation range of P-values under any sample size. Results showed that the estimation error range decreased with the increase of initial sample size, and only about 20 samples were required in each group to achieve stability (see SOM Section 4.6).

## 6. Application of P-Process to 10 P-abuse Problems

We found that P-abuse problems become easy to understand when we move up the P-value to the P-Process. We have reviewed 10 of the most critical P-abuse problems(Amrhein, Greenland, et al., 2019, Fidler, Burgman, et al., 2006, Goodman, 2008, Hoekstra, Finch, et al., 2006, Lena and Hipp, 2014, Schatz, Jay, et al., 2005) and have further expounded the three most common ones from the P-Process perspective, which were also highlighted in the introduction. Understanding these problems can be a good drop to avoid most of the P-abuse problems.

### 6.1 Question 1. Non-significant difference (P-value > 0.05) means no difference

This is the most serious and widespread P-abuse problem, 51% studies have misinterpreted non-significant results as “no effect/difference” according to the investigation by Valentin Amrhein et.al(Goodman, 2008). From the view of P-Process, this abuse will be easier to understand. P-value is not a constant value, but a stochastic process varying with sample size. Simply, we have two normal datasets with same variance but different mean, and the P-value (0.118 > 0.05) obtained from t test for them will most of the time be misinterpreted as there is “no difference” between the two groups. However, as the sample size increasing to an amount large enough, the P-Process reaches a “clear region” and the observed difference becomes clearer (Fig.6 A). So that, the whole P-Process should be considered in the process of exploring the difference, rather than just a single P-value from one hypothesis test. It is unwise to rely solely on a non-significant P-value of one hypothesis test to conclude “no difference between the two groups”.

**Fig. 6.**
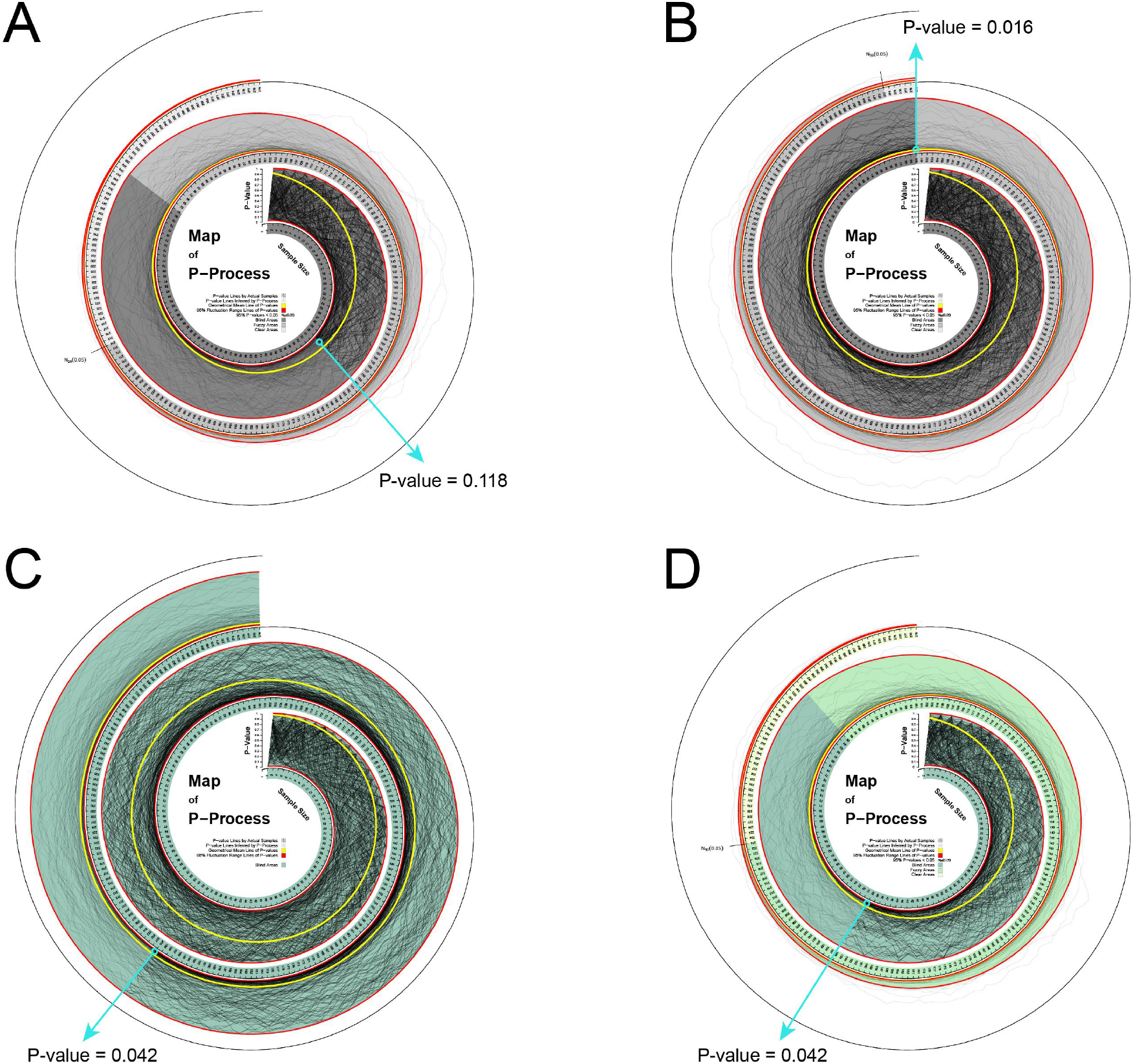
Examples of the explanation for the ten most common P-abuse problems.

### 6.2 Question 2. Significance equals to importance

From the standpoint of P-Process, P-value is not representative of the magnitude of the effect. And a significant P-value does not necessarily mean the most important difference. Considering from P-Process, an important difference should be one that requires only a small sample size to ensure 95% of the P-values less than a significance level, that is the smaller the *N*_*95*_(*α*), the more important the difference. For example, P-values obtained by dataset 1 and dataset 2 are both 0.042 (significant under the level of 0.05), but the difference of dataset 2 seems to be more important seen from the P-Process (Fig.6 C, D).

### 6.3 Question 3. P-value ≤ 0.05 and P-value > 0.05 are conflicting

This is a classic example of P-abuse problem, and largely contributes to the problem of research unrepeatability(Camerer, Dreber, et al., 2016, Camerer, Dreber, et al., 2018). From the P-Process’s view of point, P-values should not be divided into only two parts by a significance level, either black or white ones. As shown in Fig.6 A, B, although the P-values (0.118 and 0.016) are mutually contradictory under 0.05 significance level, the difference between the two sets are both really seen from P-Process. Obviously, viewed from the P-Process, there is probably no conflict between two studies with different P-values at both sides of 0.05.

In addition, we also explained the remaining seven problems from the view of P-Process, more details can be found in SOM Section 6.

## 7. Online tool ---- P-Process Map

For broader accessibility, the algorithm of P-Process was implemented into a freely available web-service (http://www.onethird-lab.com/p-process-map, SOM Section 5) that allows researchers to estimate the geometrical mean P-value as well as its fluctuation range under any sample size and to draw the P-Process Map. The only requirement for estimation is the available datasets that users could provide.

## 8. DISCUSSION

There is a widespread misconception that the P-value of an individual experiment can indicate whether a difference exists, which has largely led to an increase in non-reproducible results and the under-reporting of non-significant results. As written by Steven Goodman(Goodman, 2008), it is not accurate enough that “statistically significant results imply significant findings” or “statistically insignificant results imply no differences”.

A novel concept, the P-Process, was introduced in this article where we have evolved the P-value from a one-dimensional value to a two-dimensional stochastic process. However, the P-Process is not designed to negate P-values, but to help researchers better understand what P-value means and thus avoid their abuses. It is important to step out of the inherent thinking of local P-values and identify differences from a global perspective.

The focus of this research is on studies where the differences are not significant. By the P-Process Map showing you the whole process of P-value fluctuating, it tells the researchers why they are not significant and tells them when the difference will be significant. But when we meet the situation where the two datasets under study are from the same population, no convergence of the P-values will occur despite increasing the sample size to infinity. When this limit state occurs, we regard that there is no difference.

P-value varies with sample size, which is an unstable indicator for study results. The P-Process can effectively avoid the instability of P-values. Therefore, we recommend researchers to provide a P-Process Map in addition to the P-value given.

## Acknowledgements

YSJ conceived and contributed the work. JX, SYW, JXT, CS performed research, drafted and modified the manuscript. All authors contributed to discussing and revising the manuscript. The EWAS Project provided data support (http://www.ewas.org.cn or http://www.bioapp.org/ewasdb).

## Supplementary of Materials

### 1 Overview

In this supplementary, we achieved the following:

❖ We discussed the basic steps of hypothesis testing, and the “either-or-not” abuse of P-values. (Section 2)
❖ We defined P-Process, -ln(P-Process). (Section 4)
❖ We presented our algorithm for approximating P-Process, geometrical mean P-value and its 95% fluctuation range. (Section 4)
❖ We gave instructions on how to use the online tool, **P-Process Map**. (Section 5)
❖ We elaborated on 10 of the most common P-abuse problems from P-Process’s perspective. (Section 6)

### 2 The logic of hypothesis testing

The basic steps of the hypothesis testing are as follows(Goodman, 1993):

1. Firstly, present the null hypothesis (*H*0), no difference. Correspondingly, the alternative hypothesis is (*H*1), with difference;
2. Secondly, specify a significant level: α;
3. Thirdly, calculate the statistics through a hypothesis test method (e.g. t test, wilcoxon test, chi-square test etc.);
4. Fourthly, calculate the P-value(Wasserstein and Lazar, 2016) based on the background distribution of the statistic corresponding to the chosen hypothesis test method (e.g. t, T, χ^2^ etc.);
5. Finally, draw a conclusion based on the significant level α (i.e., *P* < α: reject *H*0, accept *H*1; *P* ≥ α: **do not** reject *H*0).

“Do not reject *H*0” is an extremely wise idea, and successfully reconciles the contradiction between the objective reality of difference and limited human cognitive ability. Because it does not force people to understand the differences that they cannot recognize, nor does it admit that there is no difference in order to comply with people’s limited cognition. “Do not reject *H*0” is not equal to “accept *H*0”. It just reminds researchers not to rush to conclusion, but to supplement the evidence and discuss or understand the difference later (Fig. S1).

**Fig. S1.**
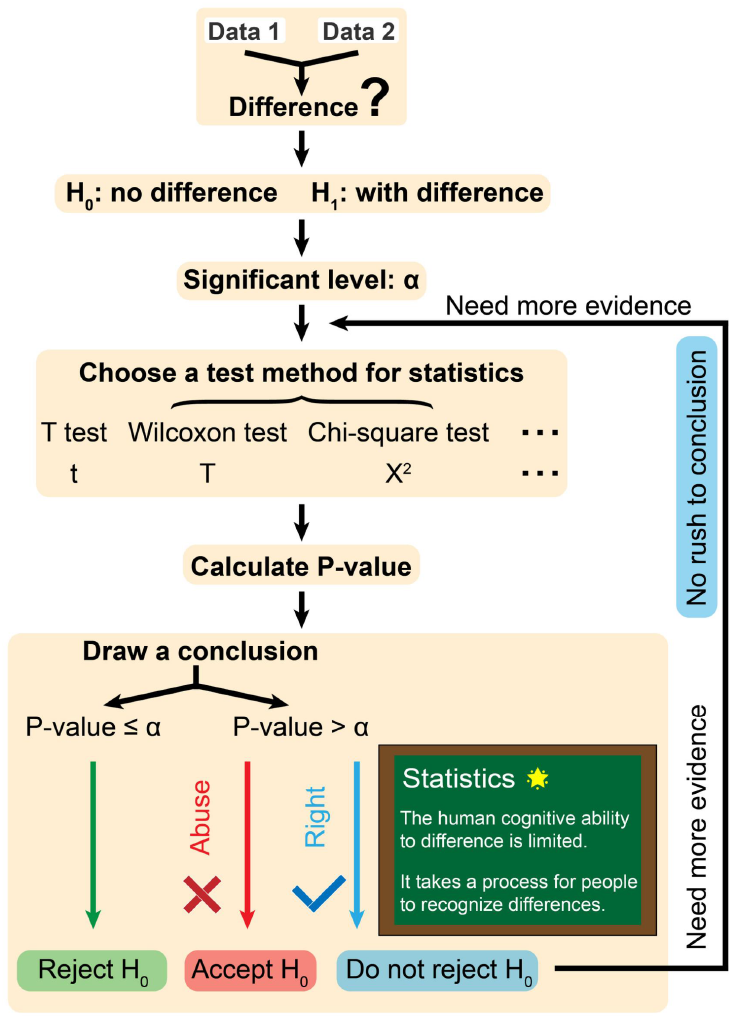
The logic of hypothesis testing.

### 3 P-value decaying process in other hypothesis test methods

There are many common hypothesis test methods for difference analysis between two groups, such as t test(Baldi and Long, 2001), wilcoxon test(Hartmann, 1978), chi-square test(Azen, Roy, et al., 1977), etc. In fact, these hypothesis test methods are essentially the same, and the main purpose of them is to identify the differences between two groups of data. The difference between them is that (i) t test identifies the difference through the mean between the two groups of data; (ii) wilcoxon test identifies the difference through the orders of data in each group; (iii) chi-square test discretizes the data and identifies the difference by comparing the frequencies between the groups.

We calculated the P-values under different sample sizes using these common hypothesis test methods separately. For t test, two normally distributed datasets were used for test (Fig. S2 A). For wilcoxon test, the test was carried on difference analysis for (i) two normally distributed datasets (Fig. S2 B), (ii) two exponentially distributed datasets (Fig. S2 C), and (iii) a normally distributed dataset and an exponentially distributed dataset (Fig. S2 D). For chi-square test, the grouped genotype (AA, AG, GG) data(Edgar, Domrachev, et al., 2002) for SNP (single nucleotide polymorphism) “rs798149” was used for test (Fig. S2 E).

All the test results showed a similar phenomenon, where in most cases the P-value decreases as the sample size increases(Demidenko, 2016) (Fig. S2).

**Fig. S2.**
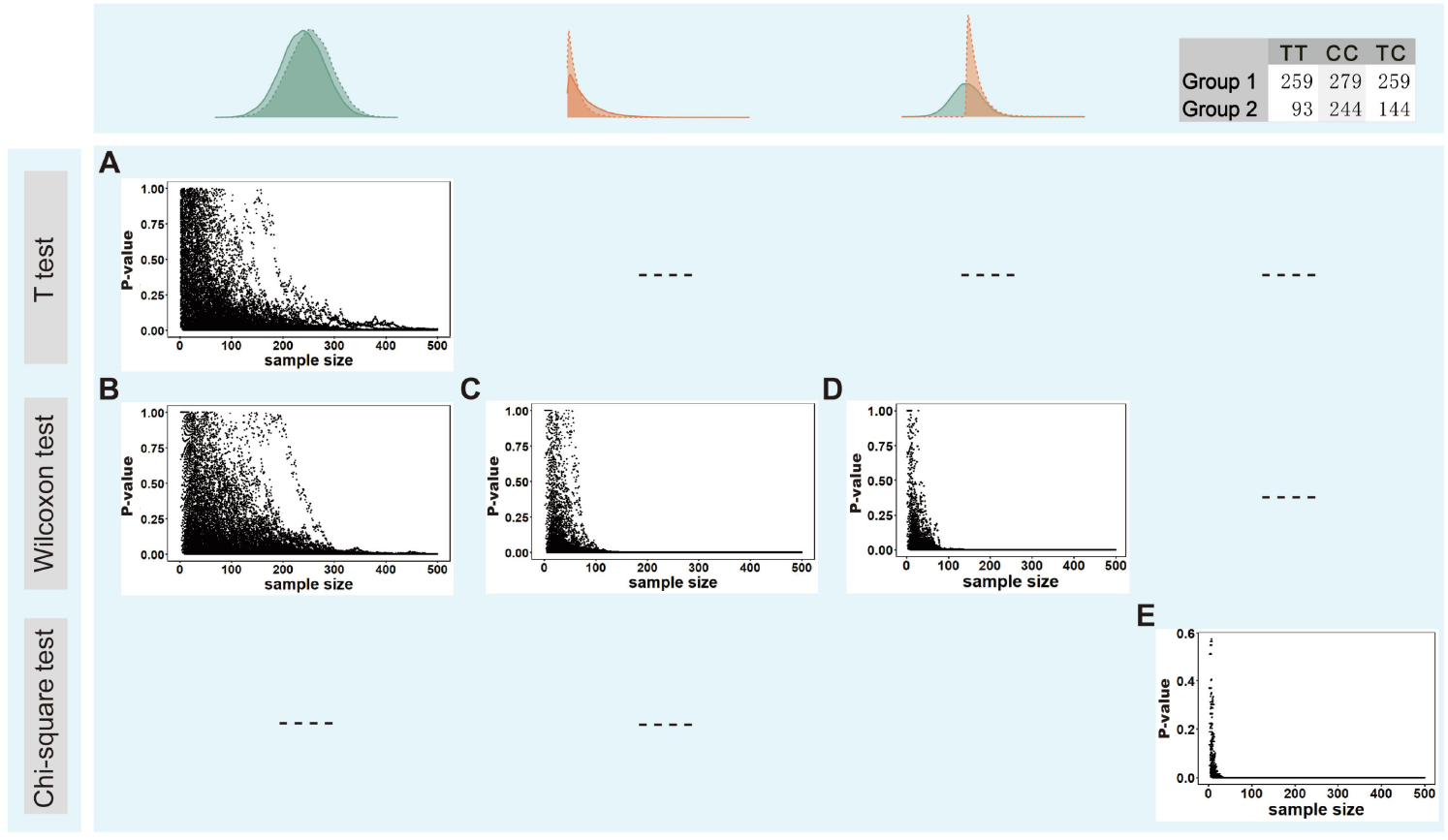
Scatter plot for P-values of **(A)** t test, **(B-D)** wilcoxon test and **(E)** chi-square test. The sample size was explored from 2 to 500. The process was randomly repeated for 100 times.

### 4 Material and Methods

#### 4.1 Main definitions

##### Definition 1

The P-Process for two given datasets A and B, which is also called Difference Recognition Process (DRP) is defined as

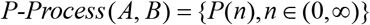

where *P*(*n*) is a random variable that denotes the P-value of hypothesis test for datasets A and B. There are five paths of P-Process for two normally distributed populations *N* (0, 3) and *N* (1, 3) which corresponds to the sample size 2 to 250 (Fig.1). The generation algorithm is described in Section 4.2.

##### Definition 2

The negative natural logarithm of P-Process for two given datasets A and B is defined as {*X*(*n*), *n* ∈ (*0*, ∞)}, and can be calculated as

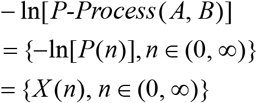

##### Definition 3

The scale uniformed map of *X*(*n*) is described as *X* ^*unif*^ (*n*). Let coordinates (*n*, − ln(*P-value*)) as the point before transformation, then the coordinates after scale unification can be calculated as (*n*, − ln(*P-value*)/*μ*), where *μ* is the average one step increment of *X*(*n*) (See Section 4.3).

#### 4.2 The algorithm for generating P-Process

In this section, we described our algorithm for generating P-Process. For two given datasets A and B, we began by outlining an idealized algorithm (Algorithm 1) for generating one path for the P-Process. For more paths of P-Process, the P-Path function was operated for several times. This was carried out by the *P-Process* function, described in Algorithm 2.

##### Algorithm 1 *P-Path*(A, B, Nall, method) #Construct the P-Process of A and B using the algorithm below.

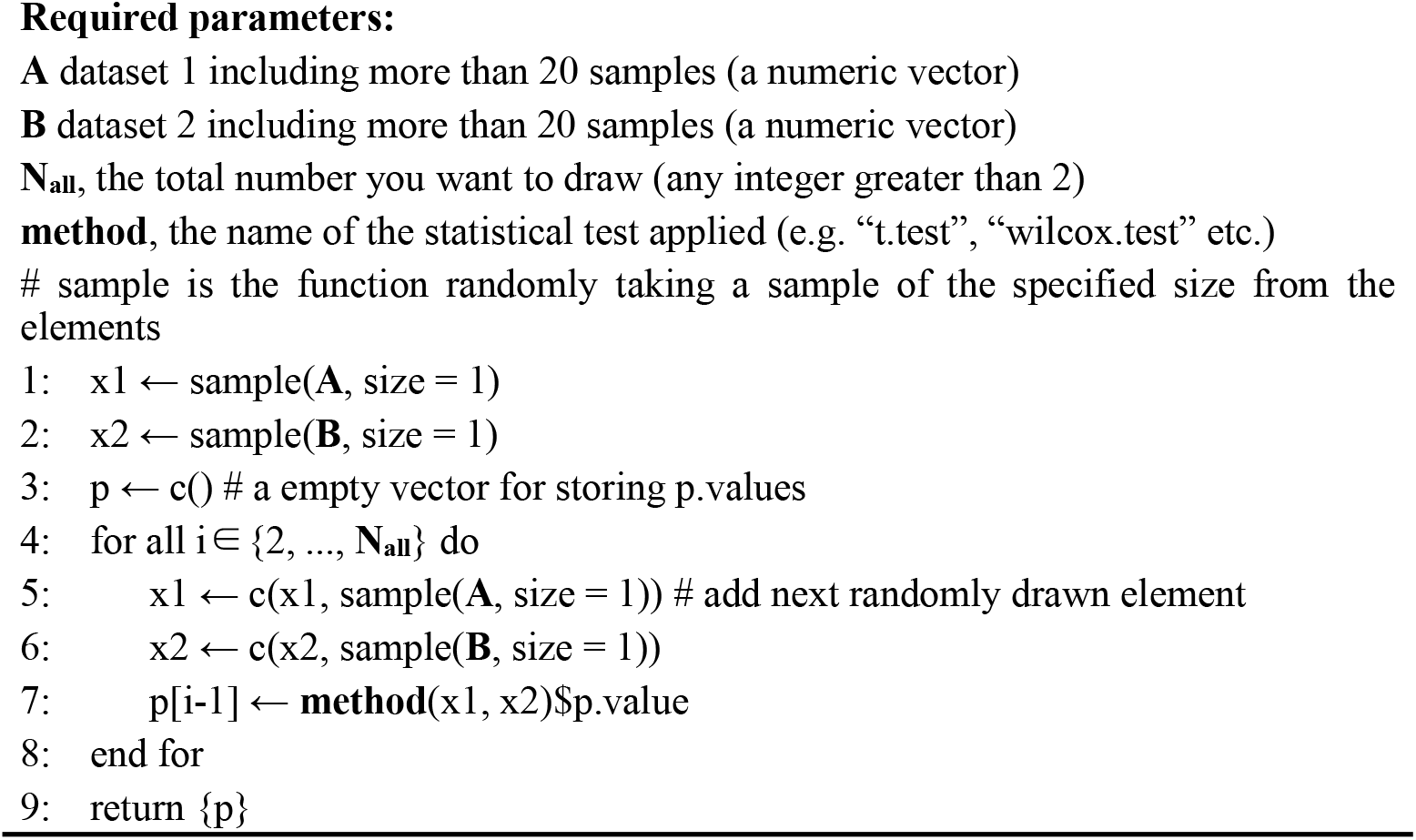

##### Algorithm 2 *P-Process*(A, B, Nall, method, Npath)

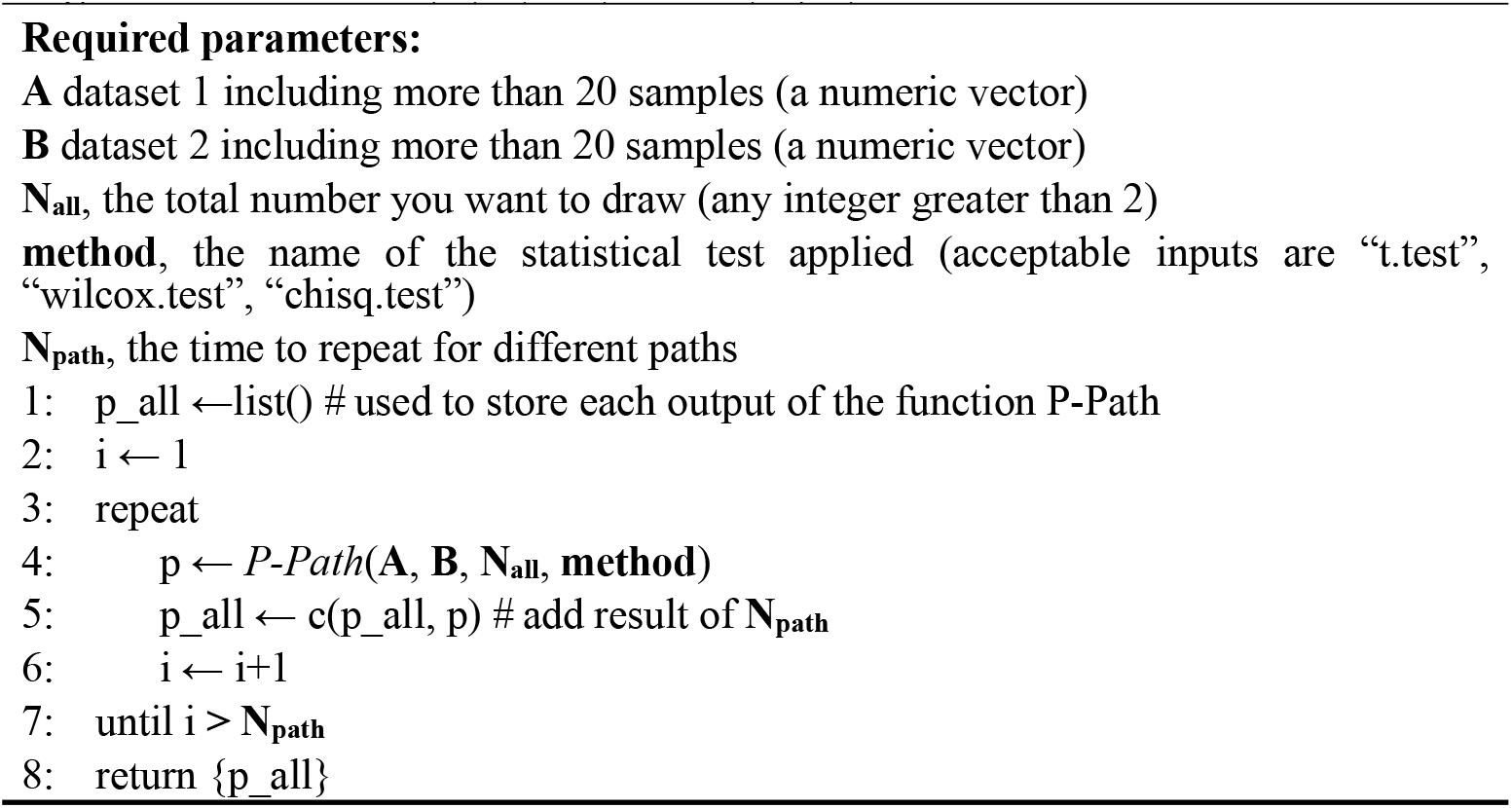

#### 4.3 The approximation for E_X_(n)

In this section, we described our algorithm for approximating the geometrical mean of P-Process, *P*(*n*). Given two datasets, we firstly obtained the *P*(*n*) (see Section 4.2) and further for *X*(*n*) (see Definition 4.2), then calculated the estimation of mean for *X*(*n*), described as a simple linear function (Fig.2 B, C):

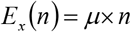

where n was the specific sample size, and *μ* was the average one step increment of *X*(*n*), which was calculated as follows:

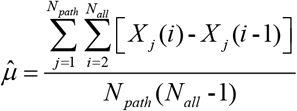

where *N* _*path*_ and *N*_*all*_ could be customized by users while generating paths of P-Process (see Section 4.2). *X* _*j*_(*i*) represented the -ln(P-value) under sample size i for the j-th path. The more obvious the difference, the larger the *μ*.

#### 4.4 The approximation for G_P_(n)

The geometrical mean of P-Process was obtained based on *E*_*X*_(*n*), and calculated as (Fig.2 D, E):

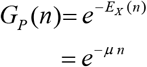

#### 4.5 The approximation for 95% fluctuation range of P-Process

##### 4.5.1 The generation of Wiener Process for estimation

We surprisingly found that the image of *X* ^*unif*^ (*n*) after 45 degrees clockwise rotation has similar morphology with Wiener Process whose one step increment is in distribution *N*(0, σ2) (Fig. S3 D), where σ is the standard variation of one step increment for *X* ^*unif*^ (*n*). We cannot help but wonder whether the fluctuation range of *X* ^*unif*^ (*n*) can be estimated with that of Wiener Process. However, there was currently a problem that the fluctuation range of the Wiener Process image was obviously a bit wider than that of *X* ^*unif*^ (*n*) after rotation.

**Fig. S3.**
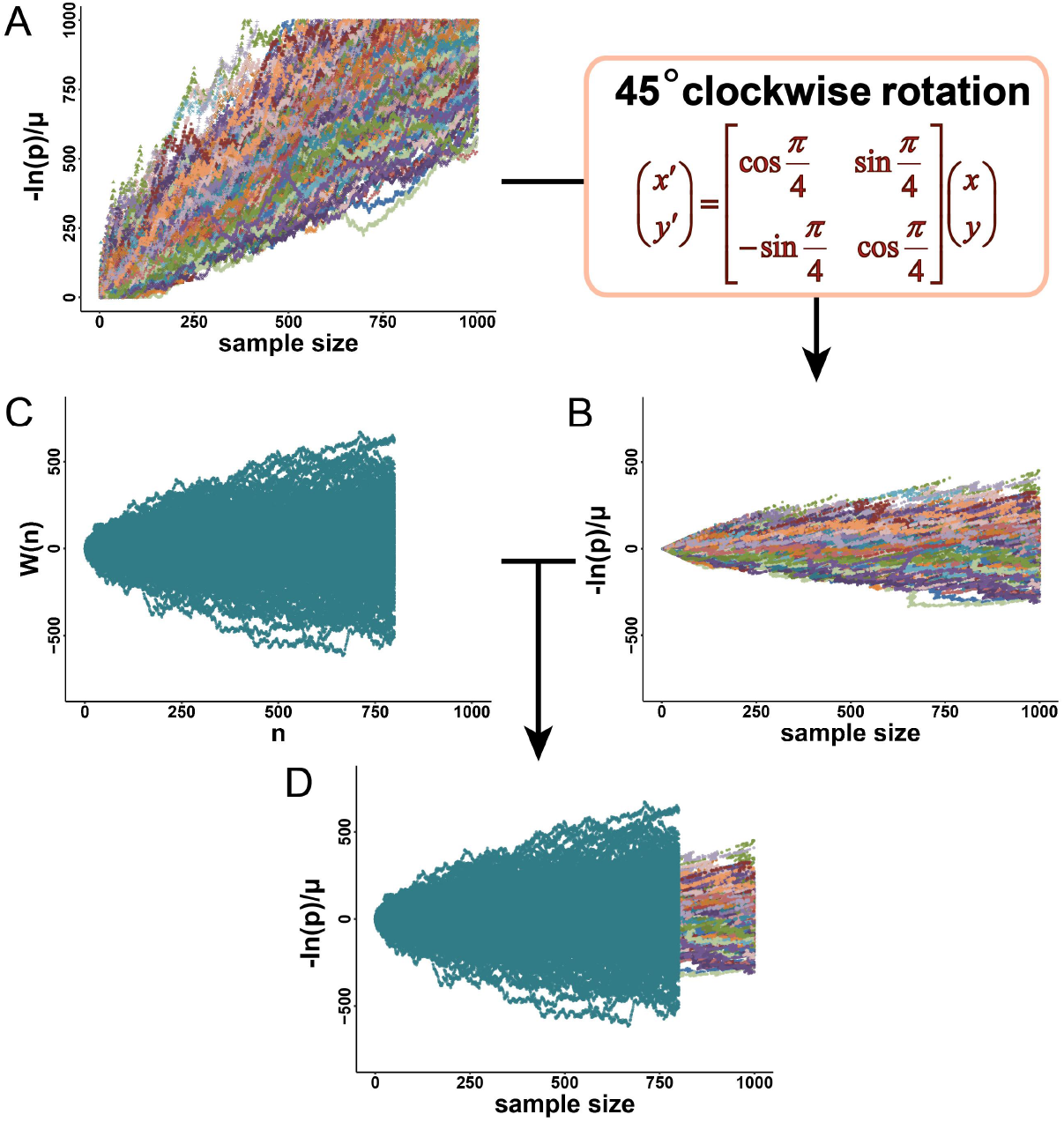
Image comparison between *X*^*unif*^(*n*) and the corresponding Wiener Process with one step increment in distribution *N*(0, σ^2^).

To improve the accuracy of estimating the fluctuation range of the P-Process, we screened k from 0.1 to 1 in steps of 0.1 and generate corresponding Wiener Process whose increment is in normal distribution *N*(0, (*k*σ)^2^) (Fig. S4). For each Wiener Process generated, the image similarity *S* _ *Score*_*k*_ was obtained by the following algorithm (Algorithm 3). And then the one with the highest similarity score was selected for estimating the fluctuation range of *X*(*n*) (Fig. S4 K).

**Fig. S4.**
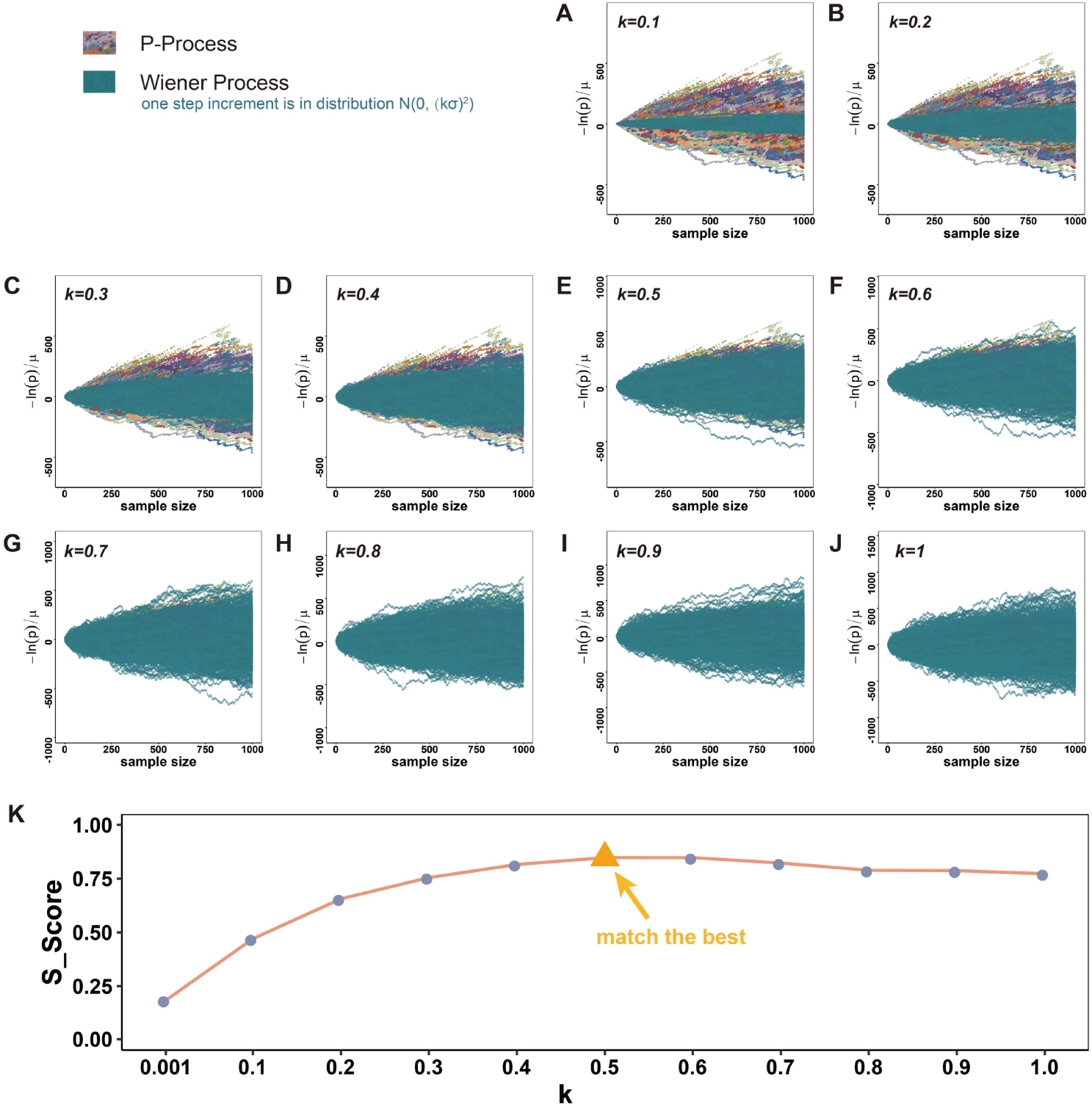
**(A-J)** An example of image comparison between the corresponding Wiener Process and *X*^*unif*^(*n*) when k is traversed from 0.1 to 1 in steps of 0.1. **(K)** Image similarity *S_Score*_*k*_.

###### Algorithm 3 *S-Score*(W, X)

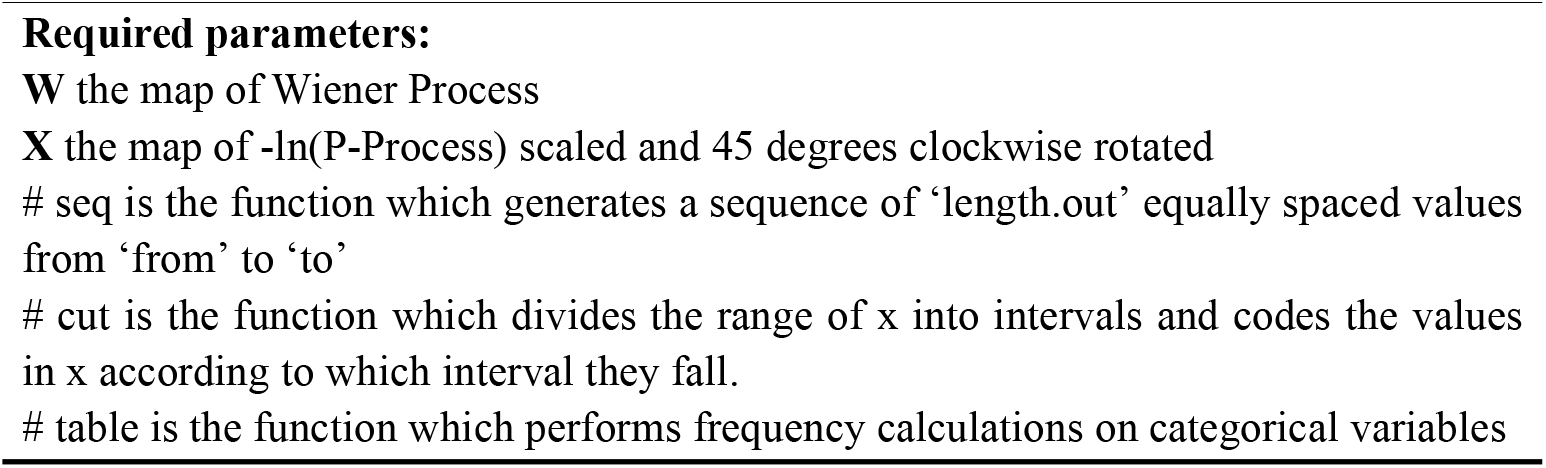
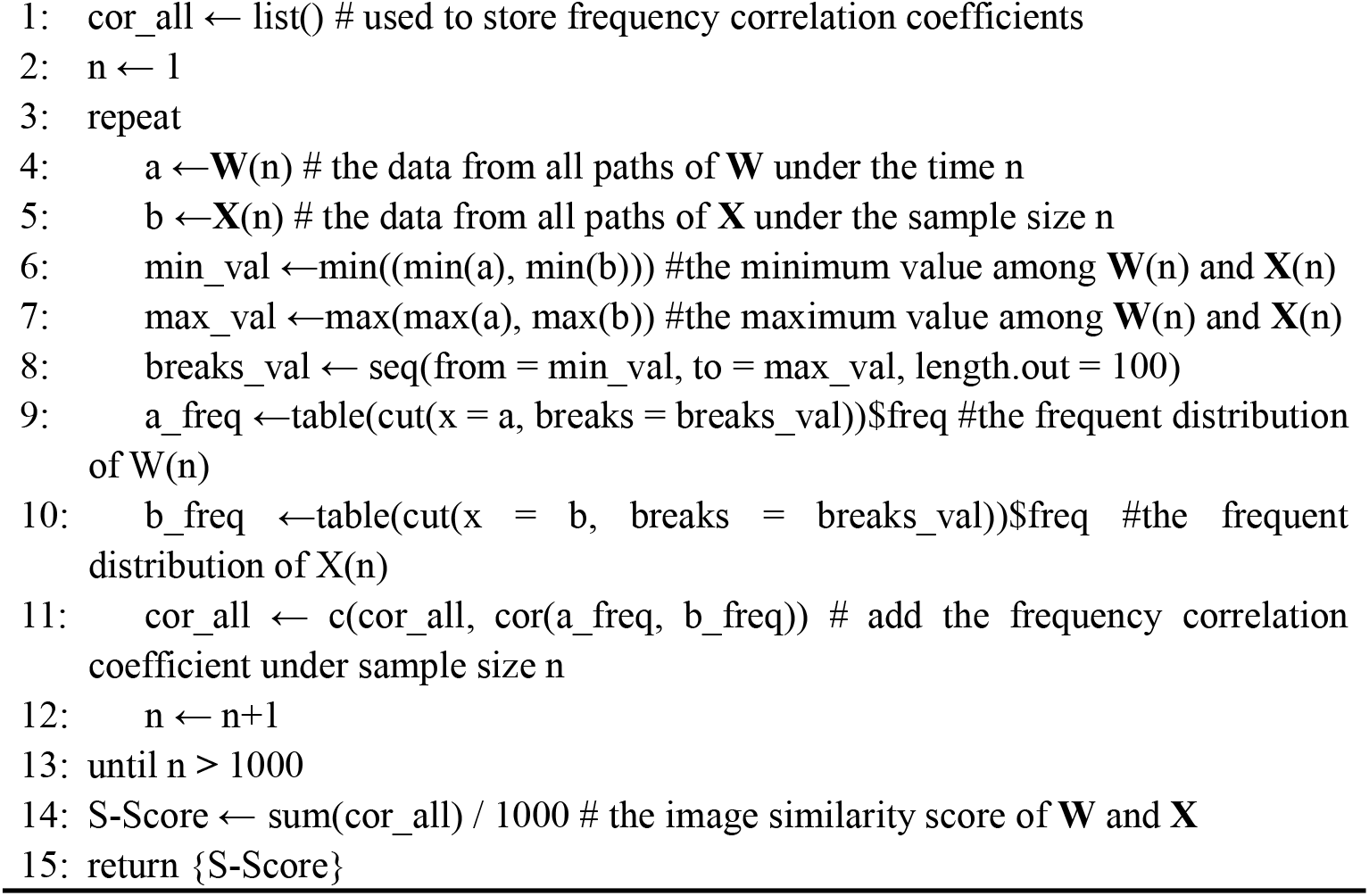

##### 4.5.2 The estimation of 95% fluctuation range of P-Process

The detailed derivation process is elaborated in the main text (Fig.3). Here is just a detailed description of the truncation value. Especially, in the derivation, the lower bound of P-values in the first small part will be negative, which is contrary to the defined range of P-values. So that we deform the derivation results and assign 0 to the part less than 0 (Fig. S5).

**Fig. S5.**
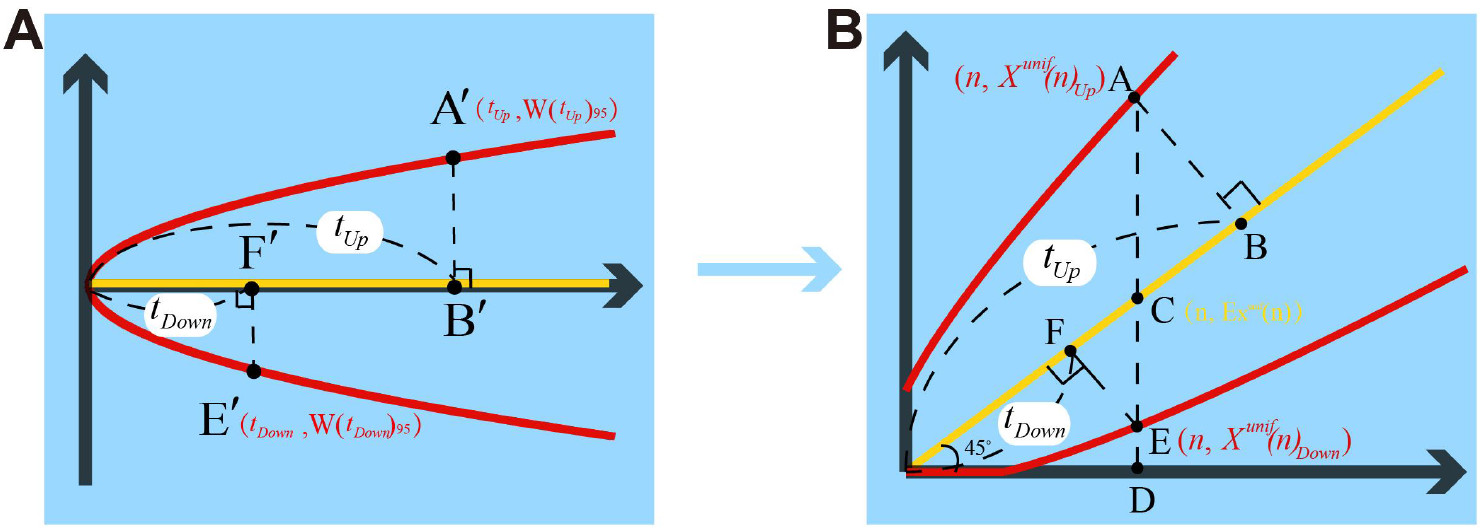
The concrete derivation process for estimating the fluctuation range of P-Process from that of Wiener Process.

##### 4.6 The performance of P-Process estimation algorithm

Here we tested the stability based on the two key parameters for the estimation of geometrical mean P-value and its 95% fluctuation range. Two normally distributed datasets *N1*(0, 3) and *N2*(1, 3) were used for testing. And the test was repeated for 40 times and the box plot was drawn (see Fig. S6). The test results showed that the estimation could be stabilized while there are only about 20 samples required.

**Fig. S6.**
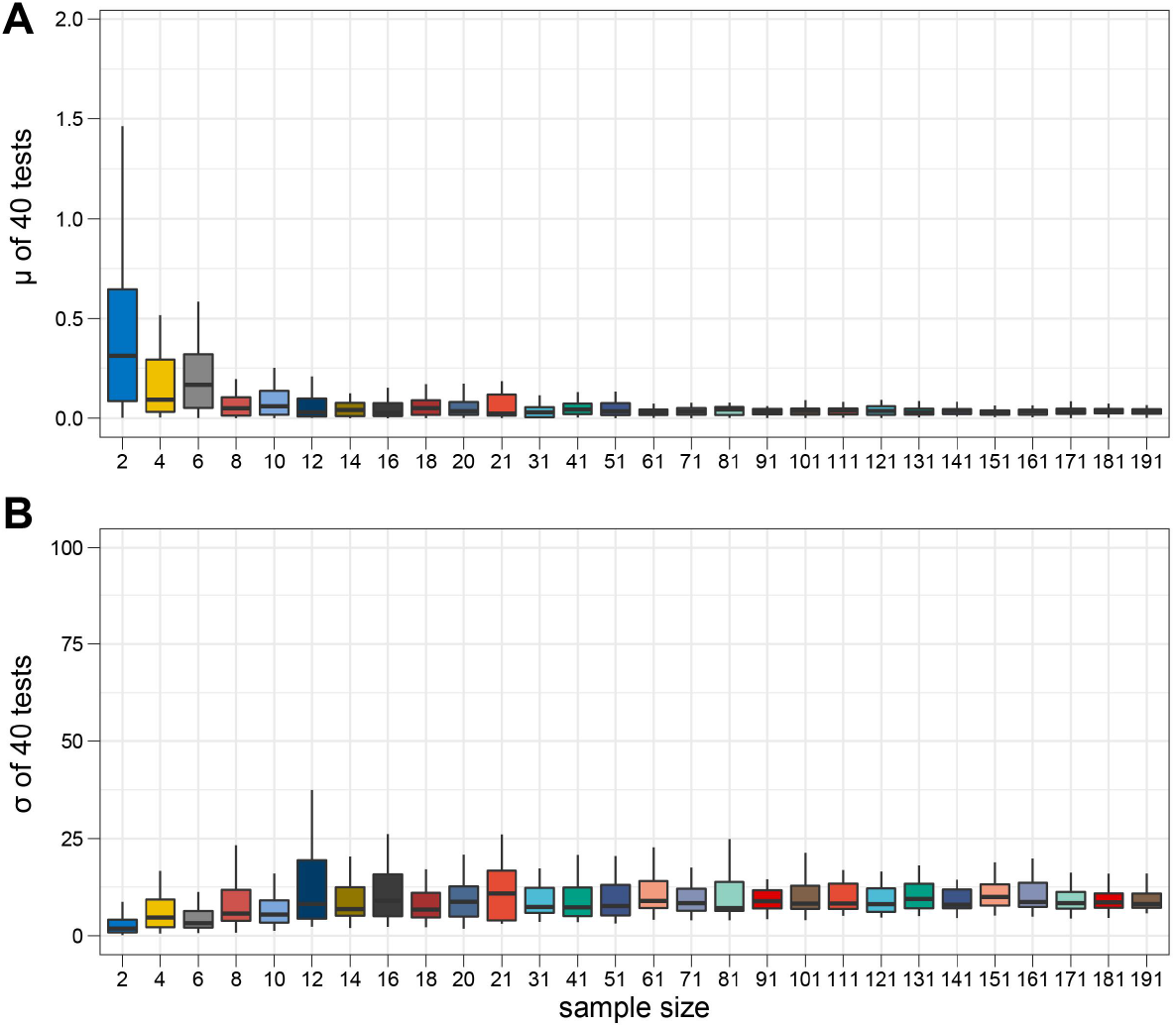
Performance testing of P-Process.

### 5 The online tool for construction of P-Process Map

For the convenience of users, we have developed an online tool **P-Process Map**, which can get the corresponding P-Process Map based on the data entered by the user. The **P-Process Map**’s user interface is easy to use and clear at a glance (Fig. S7).

**Fig. S7.**
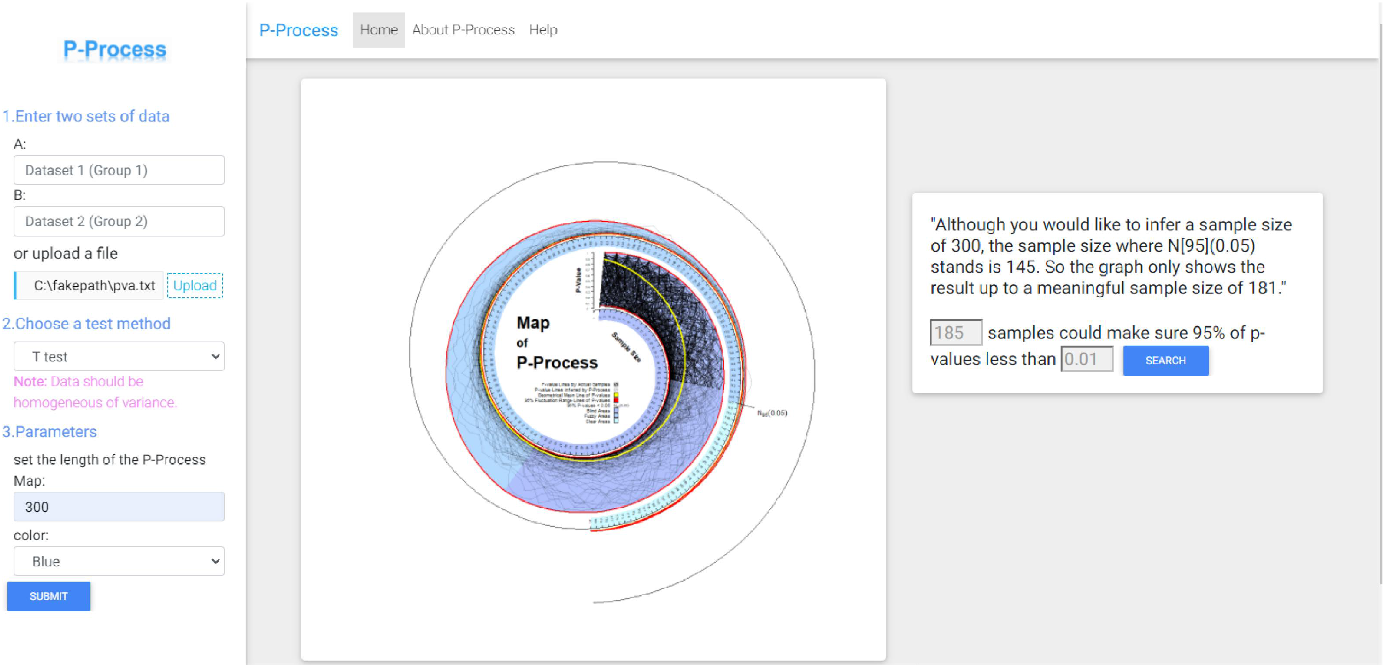
The online tool, **P-Process Map**.

Firstly, the user needs to enter (or upload with a file) two sets of data to be used for the difference identification analysis. We recommend that the sample size of the input data is preferably greater than 20 for each group. Secondly, a hypothesis test method should be chosen according to the data format entered. In general, if t test is chosen, your data should follow a normal distribution. If you choose the chi-square test, your data should be categorical data (see more details on web-page of **P-Process Map**). Thirdly, the user needs to set how long the P-Process Map is drawn, i.e. the maximum sample size will be displayed. And finally, click “Submit” to get the P-Process Map.

The trend of the P-Process can be intuitively seen in the graph at progressively increasing sample sizes. In addition, **P-Process Map** is able to give *N*_*95*_(*α*), which informs the researcher’s next study. More detailed description of the P-Process Map can be found in the section “P-Process Map” of the main text (Fig.4).

There are also some optional parameters, please see the user guide on the web page for details.

### 6 Application of P-Process to P-abuse problems

We have illustrated three of the most common problems about P-abuse in the main text from the perspective of P-Process, and found it becomes easier to understand these problems. Here we will try to describe the other seven critical P-abuse problems from the view of P-Process.

**Tab. S1.**
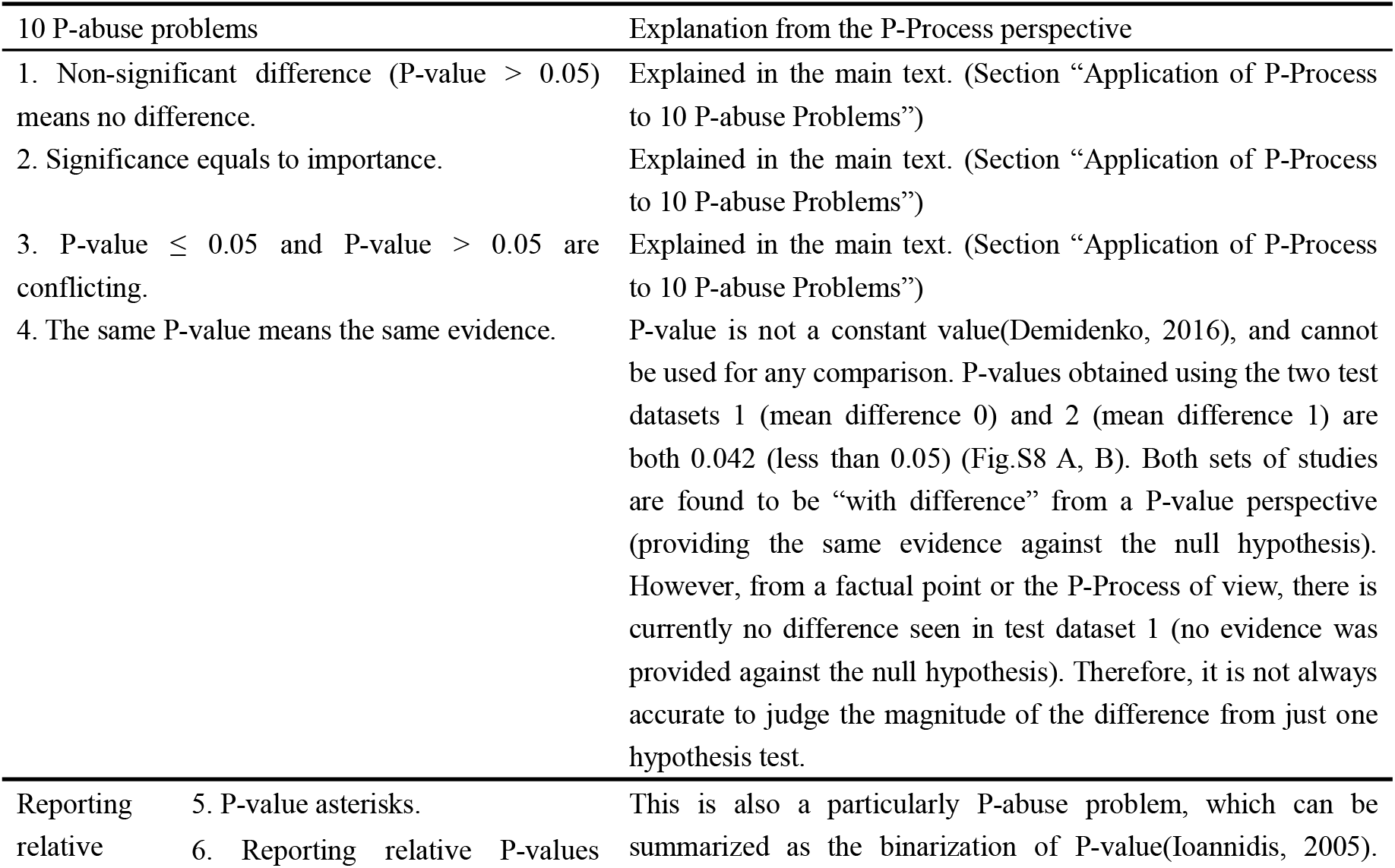
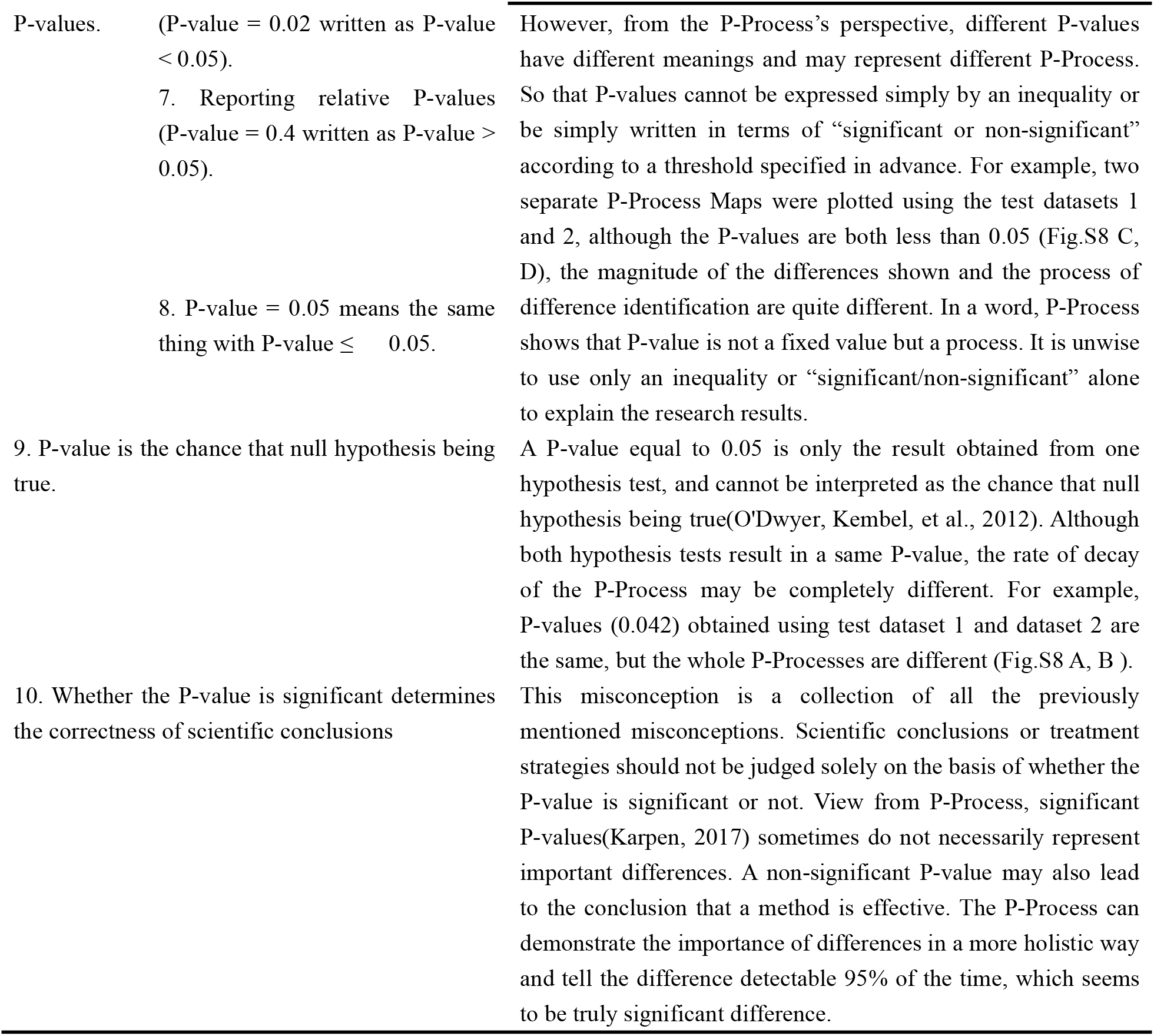
Explanation from the view of P-Process for 10 of the most common P-abuse problems.

**Fig. S8.**
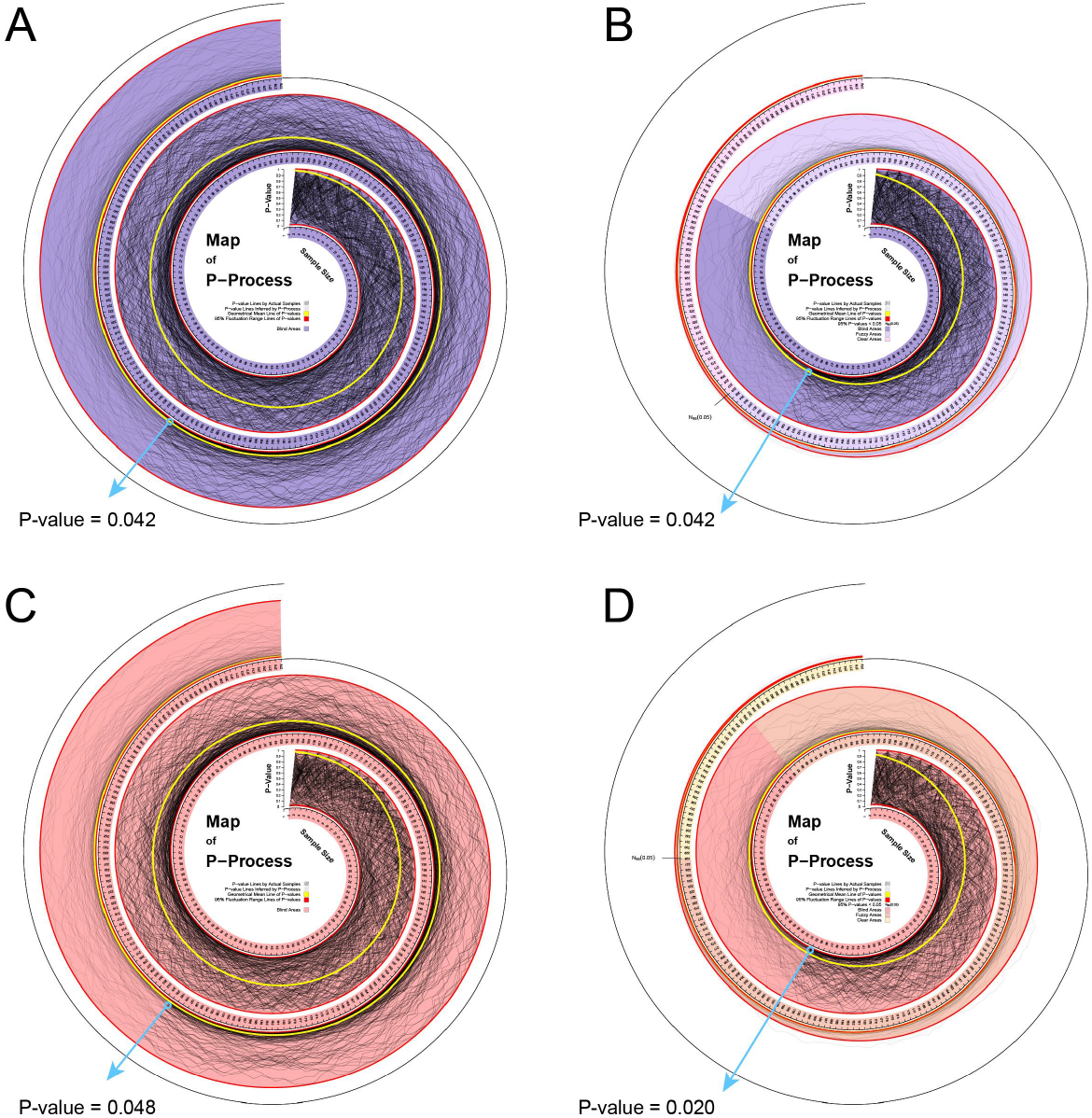
Examples for the explanation for the ten of the most common P-abuse problems.

